# Mitohormesis during advanced stages of Duchenne muscular dystrophy reveals a redox-sensitive creatine pathway that can be enhanced by the mitochondrial-targeting peptide SBT-20

**DOI:** 10.1101/2024.02.04.578832

**Authors:** Meghan C Hughes, Sofhia V Ramos, Aditya Brahmbhatt, Patrick C Turnbull, Nazari N Polidovitch, Madison C Garibotti, Uwe Schlattner, Thomas J Hawke, Jeremy A Simpson, Peter H Backx, Christopher GR Perry

## Abstract

Mitochondrial creatine kinase (mtCK) regulates the “fast” export of phosphocreatine to support cytoplasmic phosphorylation of ADP to ATP which is more rapid than direct ATP export. Such “creatine-dependent” phosphate shuttling is attenuated in several muscles, including the heart, of the D2.*mdx* mouse model of Duchenne muscular dystrophy at only 4 weeks of age. However, the degree to which creatine-dependent and -independent systems of phosphate shuttling progressively worsen or potentially adapt in a hormetic manner throughout disease progression remains unknown. Here, we performed a series of proof-of-principle investigations designed to determine how phosphate shuttling pathways worsen or adapt in later disease stages in D2.*mdx* (12 months of age). We also determined whether changes in creatine-dependent phosphate shuttling are linked to alterations in mtCK thiol redox state. In permeabilized muscle fibres prepared from cardiac left ventricles, we found that 12-month-old male D2.*mdx* mice have reduced creatine-dependent pyruvate oxidation and elevated complex I-supported H_2_O_2_ emission (mH_2_O_2_). Surprisingly, creatine-independent ADP-stimulated respiration was increased and mH_2_O_2_ was lowered suggesting that impairments in the faster mtCK-mediated phosphocreatine export system resulted in compensation of the alternative slower pathway of ATP export. The apparent impairments in mtCK-dependent bioenergetics occurred independent of mtCK protein content but were related to greater thiol oxidation of mtCK and a more oxidized cellular environment (lower GSH:GSSG). Next, we performed a proof-of-principle study to determine whether creatine-dependent bioenergetics could be enhanced through chronic administration of the mitochondrial-targeting, ROS-lowering tetrapeptide, SBT-20. We found that 12 weeks of daily treatment with SBT-20 (from day 4 to ∼12 weeks of age) increased respiration and lowered mH_2_O_2_ only in the presence of creatine in D2.*mdx* mice without affecting calcium-induced mitochondrial permeability transition activity. In summary, creatine-dependent mitochondrial bioenergetics are attenuated in older D2.*mdx* mice in relation to mtCK thiol oxidation that seem to be countered by increased creatine-independent phosphate shuttling as a unique form of mitohormesis. Separate results demonstrate that creatine-dependent bioenergetics can also be enhanced with a ROS-lowering mitochondrial-targeting peptide. These results demonstrate a specific relationship between redox stress and mitochondrial hormetic reprogramming during dystrophin deficiency with proof-of-principle evidence that creatine-dependent bioenergetics could be modified with mitochondrial-targeting small peptide therapeutics.

## Introduction

Duchenne muscular dystrophy (DMD) is a rare neuromuscular disease affecting 1 in 5,000 boys [1]. DMD is caused by X-linked recessive mutation in the dystrophin gene that almost exclusively affects males, and the loss of this structural protein triggers many cellular dysfunctions including cell membrane fragility, impaired calcium homeostasis and redox and metabolic stress [2]. Conventional glucocorticoid therapy targeting inflammation partially delays the progression of cardiac, respiratory and locomotor muscle dysfunction. However, there is no cure which underscores the need for new therapies [3]. . Furthermore, recent exon-skipping therapies, for example, have limited effects in skeletal muscle with no appreciable benefits being identified in the heart [4]. Identifying specific relationships between redox stress and metabolic dysfunction could provide foundational knowledge for pursuing new paradigms of therapy development.

A variety of mitochondrial stress responses have been reported in humans and mouse models, including elevated mitochondrial-induction of apoptosis through permeability transition pore activity, reduced oxidative phosphorylation, and elevated hydrogen peroxide emission (mH_2_O_2_) (reviewed in [5]). In cardiac left ventricle and skeletal muscle from young (4 weeks) D2.*mdx* dystrophin-deficient mice, we previously reported ADP-stimulated mitochondrial respiration was attenuated and mH_2_O_2_ was higher due specifically to a reduced ability of ADP to attenuate mH_2_O_2_ particularly when creatine was included in the experimental media compared with when creatine was absent [6, 7]. These comparisons were designed to test phosphate shuttling from mitochondrial to cytoplasmic compartments through two theoretical systems comprised of ATP export/ADP import (slow diffusion) and a faster phosphocreatine export/creatine import (faster diffusion) regulated in part by mitochondrial creatine kinase (mtCK) in the intermembrane space (reviewed in [8, 9]). Matrix ADP/ATP turnover is accelerated with creatine as mtCK activity reduces the diffusion distance for the slower diffusing ADP/ATP to the matrix-intermembrane space interface. This creatine-dependent enhancement of ADP-stimulated ATP synthesis also causes greater ADP-suppression of mH_2_O_2_ [10] by lowering membrane potential [11]. Therefore, the greater attenuations in creatine-dependent bioenergetics in 4-week-old D2.*mdx* mice suggests dystrophin deficiency impairs the more effective method of phosphate shuttling at an early stage of the disease.

These findings in 4-week-old dystrophin deficient mice provide insight into unique mitochondrial remodeling events that occur in the early stages of disease. However, the extent to which mitochondria respond in much later stages of the disease is unpredictable. While indices of mitochondrial dysfunction are to be expected, there remains the possibility of a hormetic response over time whereby mitochondria may adapt to chronic dysfunction in specific mitochondrial pathways whilst others continue to fail. Indeed, the concept of mitochondrial hormesis predicts that chronic stress can stimulate mitochondrial adaptations that lead to improved functioning [12]. Our previous findings that creatine-dependent bioenergetics are attenuated moreso than creatine independent system at 4 weeks of age in D2.*mdx* mice raises an intriguing uncertainty over whether both systems eventually fail, or one can compensate for the other. Furthermore, the findings in D2.*mdx* mice that the faster creatine-dependent system is attenuated to a greater degree than the creatine-independent system implicates mtCK – a protein known to be inhibited by reactive oxygen species (ROS) [9] - as being particularly sensitive to the redox stress of dystrophin deficiency in the disease process. However, it remains unknown if mtCK oxidation occurs during dystrophin deficiency to explain this specific remodeling of mitochondrial creatine-dependent bioenergetics at any stage of disease.In this study, we sought to determine whether both mitochondrial creatine-dependent and -independent bioenergetics are attenuated in advanced states of disease in 12-month-old D2.*mdx* mice or if either system adapts through a form of mitohormesis [12]. Our findings highlight considerable metabolic plasticity in these systems, particularly in the left ventricle, whereby severe reductions in creatine-dependent respiration and elevations in creatine-dependent mH_2_O_2_ are seemingly countered by higher respiration and lower mH_2_O_2_ in the creatine-independent system of phosphate shuttling. This observation led us to hypothesize that elevated creatine-dependent mH_2_O_2_ oxidized mtCK to explain the apparent reduction in mitochondrial sensitivity to creatine which proved to be the case. This finding of greater cysteine oxidation of mtCK linked to a unique impairment in creatine-dependent bioenergetics led us to perform a separate investigation demonstrating proof-of-principle that cardiac creatine-dependent bioenergetics are enhanced with 12 weeks of treatment with the ROS-lowering mitochondrial-targeting tetrapeptide SBT-20 [13].

## Methods

Male D2.*mdx* mice originated from breeding colonies maintained at York University (Toronto, Canada) and sourced from The Jackson Laboratory (Bar Harbor, United States). DBA/2J wild type were purchased from The Jackson Laboratory at 4-5 weeks of age and aged in-house.

In the first part of this study, D2.*mdx* and wild type mice were aged to 52 weeks (12 months). Two to three days prior to tissue removal, mice were assessed for 24-hour voluntary wheel running, hang time using an inverted cage lid and forelimb grip strength. Mice also received a single micro computed tomography scan for measurement of lower limb muscle volume. In the second part of the study, beginning at 4-days of age, D2.*mdx* mice received subcutaneous injections of 5mg/kg SBT-20 (Stealth Biotherapeutics; Newton, MA, USA) 7 days/week continuously for 12 weeks. Thereafter, ultrasound assessments of cardiac function were performed, and muscles were removed.

Detailed methodology, including information on permeabilized muscle fibre bundles, high resolution respirometry, mitochondrial H_2_O_2_ emission, calcium retention capacity assessments, glutathione, western blots, redox assessments of mtCK, and statistical methods, are described in detail in the *Supplemental Information (**Appendix A**)*.

## Results

### Mitochondrial respiration and mH_2_O_2_: modeling metabolic demand and phosphate shuttling

At 12 months of age, D2.*mdx* mice showed reduced body weight, lower limb muscle volume and cage hang time compared with age matched DBA/2J wild type mice (**Figure 1A-E**) as expected with this model [7]

**Figure 1.**
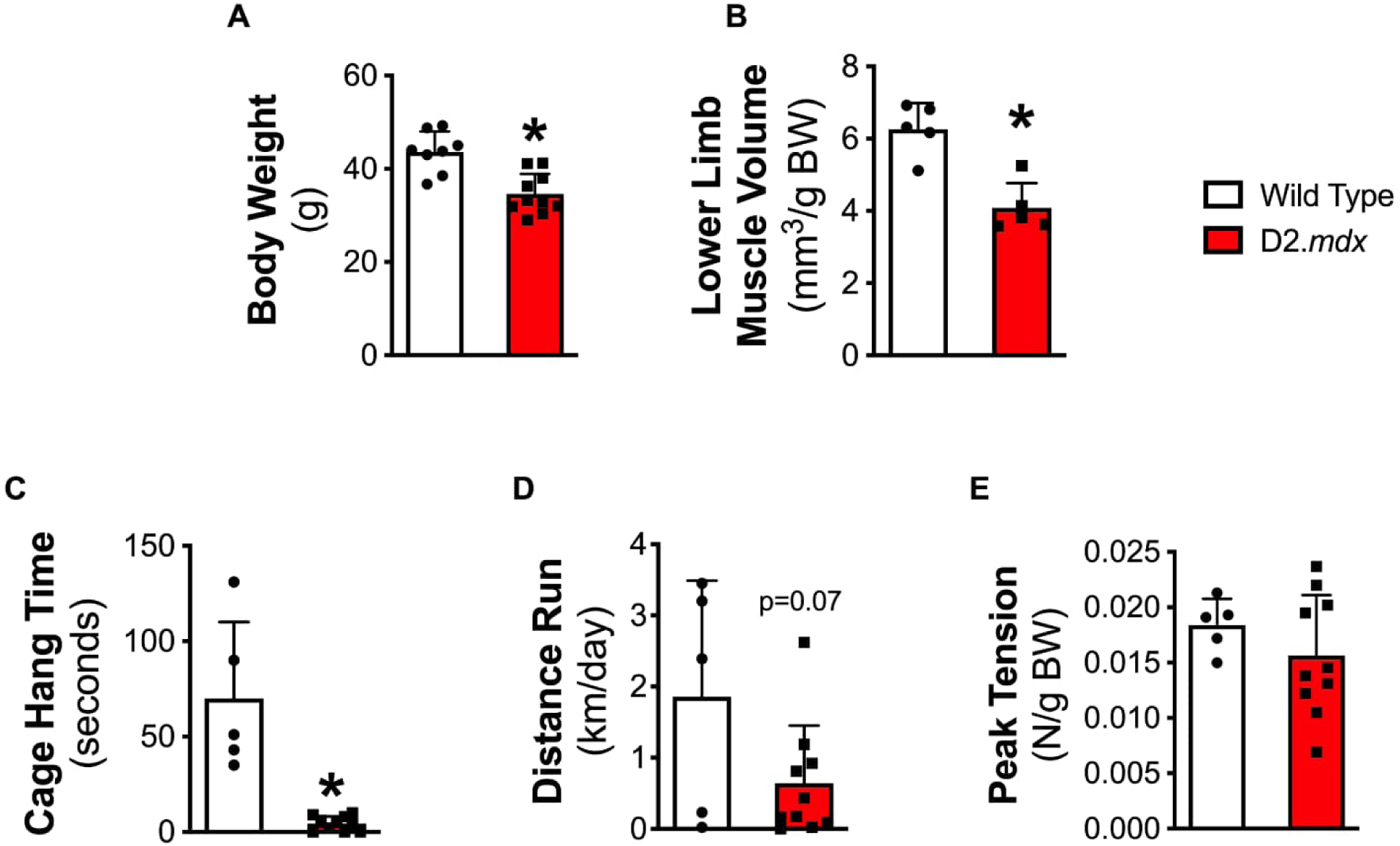
Anthropometrics and functional testing in 12-month-old D2.mdx mice. Body weight (**A**, n=8-10), lower limb muscle volume assessed by microCT (**B**, n=5), cage hang time (**C**, n=5-10), voluntary wheel running in 24 hours (**D**, n=5-9) and grip strength (**E**, n=5-10). Data were analyzed by unpaired t-tests. Results represent mean +/- SD; *p<0.05 compared with wild type.

Creatine-dependent versus creatine-independent regulation of ADP-stimulated respiration were assessed by placing separate permeabilized left ventricle fibre bundles in experimental media with 20mM creatine to saturate mtCK [14, 15] or in media without creatine to model both theoretical models of phosphate shuttling (**Figure 2A and B**). We found that left ventricles from 12-month-old D2.*mdx* mice have lower creatine-dependent Complex I-supported respiration compared to wildtype when stimulated by both low and high [ADP] modeling a range of metabolic demands for mitochondrial ATP synthesis (**Figure 2C and D**). D2.*mdx* surprisingly showed increased respiration compared to wildtype when assessed in the absence of creatine at both low and high [ADP] suggesting the slower but direct ADP/ATP cycling system was upregulated despite reductions in the faster creatine-dependent system. Furthermore, the higher respiration seen in the condition with creatine compared to the absence of creatine in wildtype (**Figure 2C and D**) is consistent with the known stimulatory effects of creatine on enhancing ADP-stimulated respiration reflective of faster adenylate cycling (**Figure 2A**). In this regard, a critical observation is that this stimulatory effect of creatine was lost in the 12-month D2.*mdx* mice as shown by similar respiration rates when creatine was either present or absent (**Figure 2C and D)** at both low or high [ADP]. Collectively, these results demonstrate a loss of mitochondrial creatine sensitivity in 12-month-old D2.*mdx* mice as well as an apparent compensation in the slower creatine-independent model of adenylate cycling.

**Figure 2.**
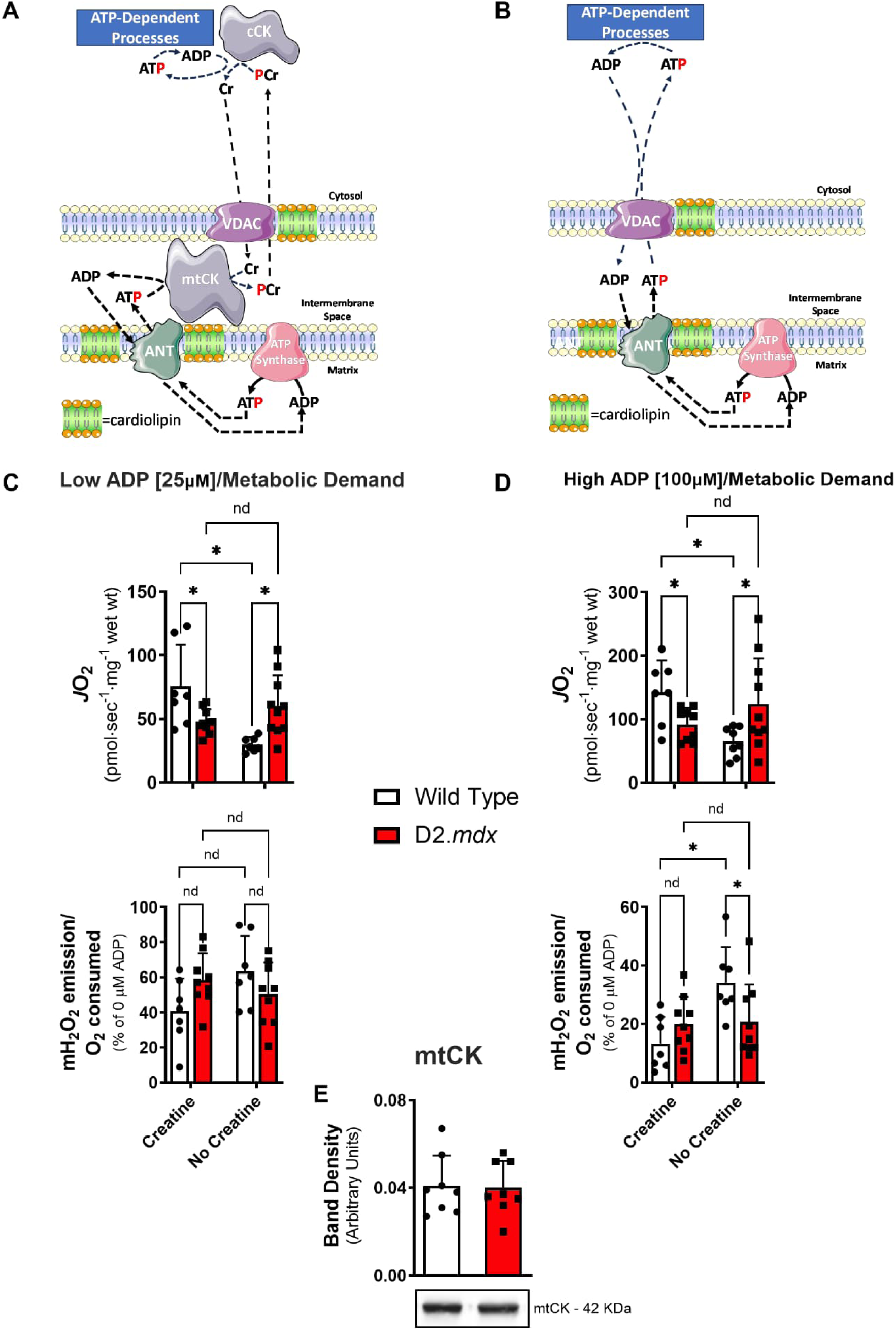
Complex I-supported mitochondrial respiration and mitochondrial H_2_O_2_ emission (mH_2_O_2_) in left ventricles from 12-month-old D2.mdx mice. Creatine-dependent (**A**) and - independent (**B**) ADP-stimulated respiration (JO_2_) and ADP-suppression of mH_2_O_2_ during the process of oxidative phosphorylation (mH_2_O_2_/O_2_) with low [ADP] (25uM; **C**) and high [ADP] (500uM; **D**) were assessed in cardiac left ventricle permeabilized muscle fibre bundles by stimulation with pyruvate (5mM respiration, 10mM mH_2_O_2_) and malate (2mM) to stimulate Complex I with NADH. Mitochondrial creatine kinase (mtCK) total lysate protein content were assessed in heart samples remaining after removal of left ventricles (**E**). mH_2_O_2_ represents forward electron transfer that was achieved with NADH generated by pyruvate and malate to stimulate complex I with and without creatine in the experimental media. mH_2_O_2_ arising from electron slip in the electron transport chain during the process of oxidative phosphorylation is positively associated with membrane potential is therefore suppressed by [ADP] [11]. For A and B, several decades of research has contributed to this theoretical model whereby the matrix-derived ATP is transported through the inner membrane transporter ANT (adenine nucleotide translocase) to the intermembrane space (see [19] for review). In the presence of creatine, a phosphate is transferred from ATP to creatine by mtCK to produce phosphocreatine (PCr). The ADP product cycles back to the matrix while PCr is exported to the cytoplasm (theorized through VDAC; voltage dependent anion carrier) where it is used by cytoplasmic creatine kinase (cCK) to re-phosphorylate ADP to ATP to support ATP-dependent proteins with creatine returning to the mitochondria. This PCr/creatine system cycles faster than ATP/ADP due to faster diffusion kinetics of both PCr and creatine relative to ADP and ADP, and is estimated to represent up to 80% of phosphate exchange between mitochondria and cytoplasmic compartments vs 20% for the direct ATP/ADP shuttle. mtCK, ANT and VDAC are thought to be bound to cardiolipin (see [9] for review). Both systems are thought to be active in vitro in the presence of creatine. Diffusion distances are not to scale. Data were analyzed by Two-way ANOVA for data in panels C and D, and unpaired t-test for panel E. Results represent means +/- SD; n=7-10. *p<0.05; nd means ‘no difference’.

In the presence of creatine, mH_2_O_2_ driven by Complex I-supported mH_2_O_2_ (NADH) through forward electron transfer was not different between wildtype and D2.*mdx* at low and high [ADP] (**Figure 2C and D**) but was higher in D2.*mdx* with reverse electron transfer with high [ADP] (**Supplemental Figure S1A and B**). This was noted when expressing mH_2_O_2_ relative to oxygen consumption at the same ADP concentration made in parallel fibre bundles to gain insight into how mH_2_O_2_ is regulated during the process of oxidative phosphorylation. Separate post-hoc analyses examining the ability of creatine to attenuate mH_2_O_2_ [16] revealed that wildtype hearts showed the expected effect whereby mH_2_O_2_ was lower when creatine was present compared to its absence when stimulated with both forward electron transfer during high [ADP] (**Figure 2C and D**) and reverse electron transfer during both low and high [ADP] (**Supplemental Figure S1A and B**), but this effect was lost in the D2.*mdx* whereby mH_2_O_2_ was similar in the presence and absence of creatine (**Figure 2C and D, Supplemental Figure S1A and B**). Given creatine is known to accelerate ADP/ATP cycling which enhances the well-established effect of ADP in lowering membrane potential-dependent mH_2_O_2_ ([6, 16–18]; see discussion), these findings suggest that creatine is less capable of stimulating respiration, as described above, and attenuating mH_2_O_2_ in the left ventricles of 12-month-old D2.*mdx* mice. Furthermore, similar to the findings with respiration, there was an apparent compensation in the creatine-independent system whereby mH_2_O_2_ in the absence of creatine was lower in D2.*mdx* than wildtype with high [ADP] when driven by forward electron transfer (**Figure 2D**) or with both low and high [ADP] when driven by reverse electron transfer (**Supplemental Figure S1A and B**).

There were no changes in mtCK protein content of the left ventricles (**Figure 2E)** indicating that factors independent of mtCK content may mediate the loss of creatine sensitivity in D2.*mdx* hearts.

Collectively, these results demonstrate divergent remodeling of left ventricular mitochondrial-cytoplasmic ADP/ATP cycling in 12-month-old D2.*mdx* mice whereby impairments in the faster creatine-dependent regulation of ADP-stimulated respiration and suppression of mH_2_O_2_ are countered by apparent compensations in the slower creatine-independent system.

We also questioned whether the reductions in creatine-dependent ADP governance of respiration and mH_2_O_2_ were unique to the heart or if it occurred in other muscles in D2.*mdx* at 12 months of age.. Unlike the dystrophic heart (**Figure 2C, D**), creatine stimulated increases in respiration in the diaphragm of 12-month-old D2.*mdx* but, similar to the heart, did not attenuate mH_2_O_2_ (**Supplemental Figure S2B, C, E, and F**). In the diaphragm, creatine-stimulated respiration appeared to be greater during low and high [ADP] in 12-month-old -matched wildtype vs D2.*mdx* suggesting a partial impairment in creatine sensitivity nonetheless occurs in dystrophic diaphragm. However, unlike the left ventricle, there were no apparent compensatory increases in respiration in the absence of creatine but lower mH_2_O_2_ in this condition, similar to the heart, were observed (**Supplemental Figure S2B, C, E, and F**). Also, no effect of creatine was seen in quadriceps or white gastrocnemius in 12-month-old wildtype, in contrast to the effects seen in our previous work at 4 weeks ([17, 18] see discussion), or in 12-month-old D2.*mdx* (**Figure 2C, D**).

We next explored whether changes in bioenergetics in each muscle type were related to altered contents of electron transport chain and phosphate shuttling components that are stimulated in most of the bioenergetic protocols employed in this study. In the heart, no changes in specific subunits of the electron transport chain, ANT1 or VDAC2 occurred, similar to mtCK (**Figure 2E**), suggesting changes in respiration and mH_2_O_2_ might be linked to altered intrinsic activities, but this would require further investigation. Pyruvate dehydrogenase contents or activities, which regulates NADH production by pyruvate in these protocols, were not assessed. In the diaphragm of 12-month-old D2.*mdx* mice, certain components of the electron transport chain as well as ANT1 and VDAC2 were lower (**Supplemental Figure S3B, C**) which could explain the lower respiration. In the white gastrocnemius, increased subunits of Complex I and IV as well as VDAC2 were observed (**Supplemental Figure S3B, C**). The lack of differences in respiration in the white gastrocnemius suggests the higher contents of these proteins (all of which are involved in these ADP-stimulated respiration protocols) may have offset reductions in their specific activities in order to ‘maintain’ normal respiration rates expressed per mg tissue (**Supplemental Figure S2B, C, E, and F)**. Increases in complex II in the quadriceps would not be expected to influence the Complex I-stimulated respiration or mH_2_O_2_ protocols used in this study. Collectively, there are heterogeneous mitochondrial alterations across muscle type in 12-month-old D2.*mdx* but the loss of creatine sensitivity seems to be predominant in the cardiac left ventricles despite no changes in contents of many proteins stimulated in the bioenergetic protocols used in this study.

In an effort to explain why mitochondrial creatine sensitivity was uniquely impaired in the dystrophic heart and considering there were no changes in mtCK protein content of the left ventricles (*see* **Figure 2E**), we next questioned whether mtCK was modified through redox-linked post-translational modifications given mH_2_O_2_ was elevated.

### Cellular redox state and mtCK thiol oxidation

The increased creatine-dependent mH_2_O_2_ during the process of oxidative phosphorylation in 12-month-old D2.*mdx* mice guided us to hypothesize that mtCK thiols would be more oxidized than in wild type given mtCK is a redox-sensitive protein (reviewed in [9]). We next immunoprecipitated mtCK from left ventricles (**Supplemental Figure S3D**) and incubated the extract in a maleimide-tagged fluorophore that binds irreversibly to reduced cysteine thiols, with no affinity to oxidized thiols, as previously described ([17, 20, 21], **Supplemental Figure S3**). This experiment demonstrated that mtCK is more oxidized in the left ventricle of 12-month-old D2.*mdx* mice (**Figure 3A**). This was related to a more oxidized glutathione (H_2_O_2_ scavenger) redox state as reflected by a lower GSH:GSSG (reduced to oxidized glutathione redox buffer) in lysate from left ventricles due apparently to high variability in GSSG (**Figure 3B-E**). This oxidized environment in the left ventricle (**Figure 3B-E**) was more pronounced than other specific muscles, although the diaphragm also showed a lower GSH:GSSG despite increases in both GSH and GSSG (**Supplemental Figure S3A)**. No changes were observed in quadriceps and white gastrocnemius (**Supplemental Figure S3A**).

**Figure 3.**
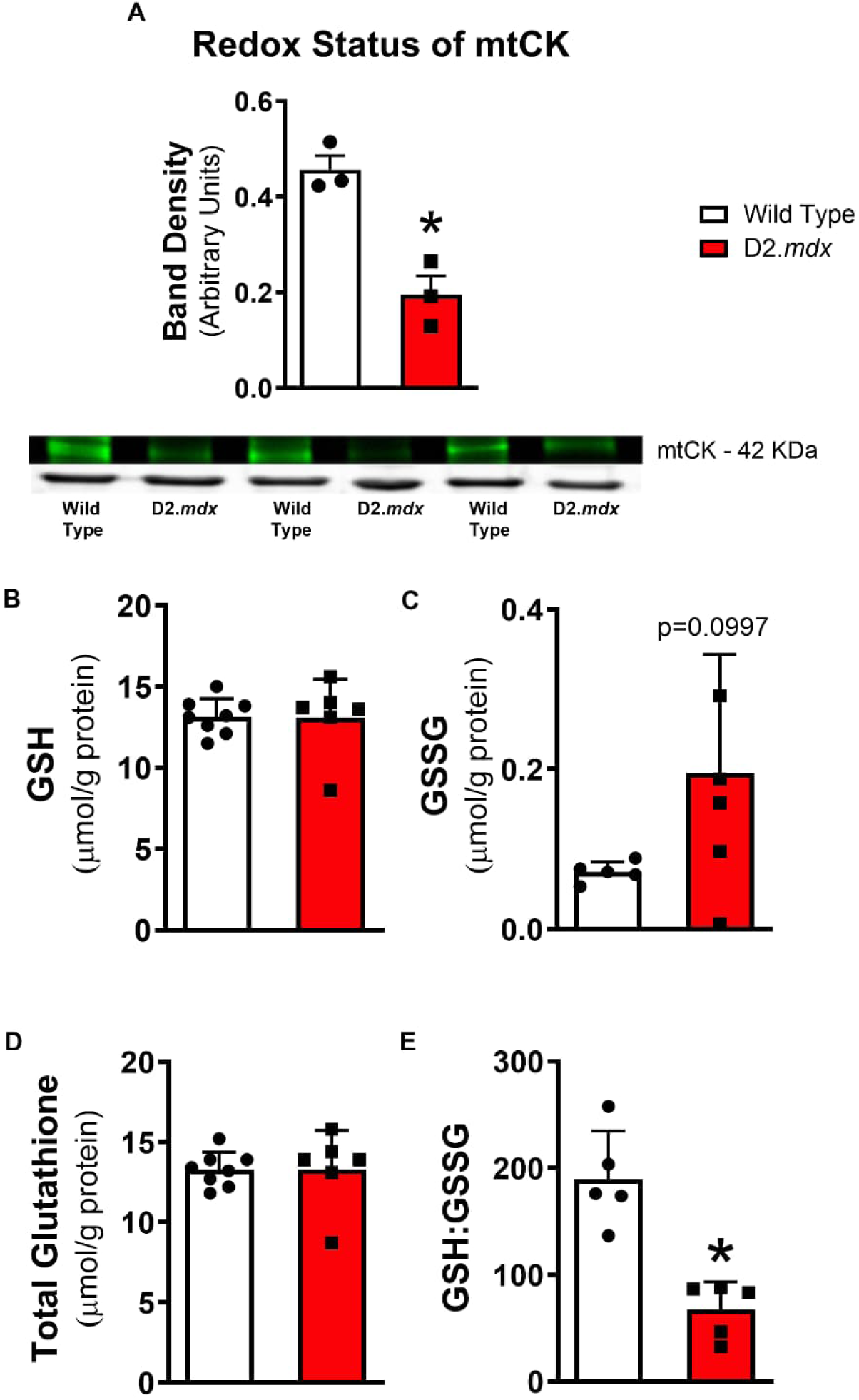
Mitochondrial creatine kinase (mtCK) cysteine redox state and cellular glutathione redox state in the heart from 12-month-old D2.mdx mice. Measurements were made in frozen heart following the removal of left ventricles. Greater cysteine oxidation on immunoprecipitated mtCK from D2.mdx left ventricles is demonstrated by lower binding of the maleimide-tagged fluorescent IR-dye 800 CW probe compared to wild type (**A**). Glutathione was measured in left ventricle lysates using HPLC-UV for the detection of GSH (**B)** and HPLC-fluorescence for GSSG (**C**). The GSH:GSSG ratio (**D**) and total glutathione (GSH + 2x GSSG; **E**) were calculated from GSH and GSSG. Data were analyzed by unpaired t-tests between wild type and D2.mdx Results represent mean +/- SD; n=4-8. *p<0.05 compared to wild type.

There were no differences in ANT1, VDAC2 protein contents or subunits of the electron transport chain in the left ventricle (**Supplemental Figure S3B and SC**). While we did not assess their post-translational modifications, these collective observations guided us to examine mtCK-linked creatine-dependent bioenergetics in the heart in more detail.

### In vivo treatment with the mitochondrial ROS-lowering peptide SBT-20

Given we observed attenuated creatine-sensitive control of bioenergetics by ADP in the left ventricle, as the second part of the study we performed a pilot study with *in vivo* injections of the mitochondrial targeting ROS-lowering peptide SBT-20 in D2.*mdx* mice from 4 days of age to ∼12.5 weeks of age. Treatment at an earlier age was chosen given a 12-month protocol was not possible. This 12-week treatment protocol increased pyruvate-supported ADP-stimulated respiration in the presence of creatine compared to saline-treated D2.*mdx* mice and had no effect on creatine-independent respiration at low or high [ADP] (**Figure 4A and C)**. Unlike 12-month-old D2.*mdx* where creatine did not increase cardiac mitochondrial respiration compared to the absence of creatine (**Figure 2C and D**), left ventricles from 12-week-old mice appeared to retain some creatine sensitivity given respiration was higher in D2.*mdx* saline in the presence of creatine vs in the absence of creatine (**Figure 4A and C)**. Nonetheless, the results demonstrate that SBT-20 has a specific action of enhancing creatine-dependent respiration in D2.*mdx* mice given no effect was seen in the creatine-independent condition.

**Figure 4.**
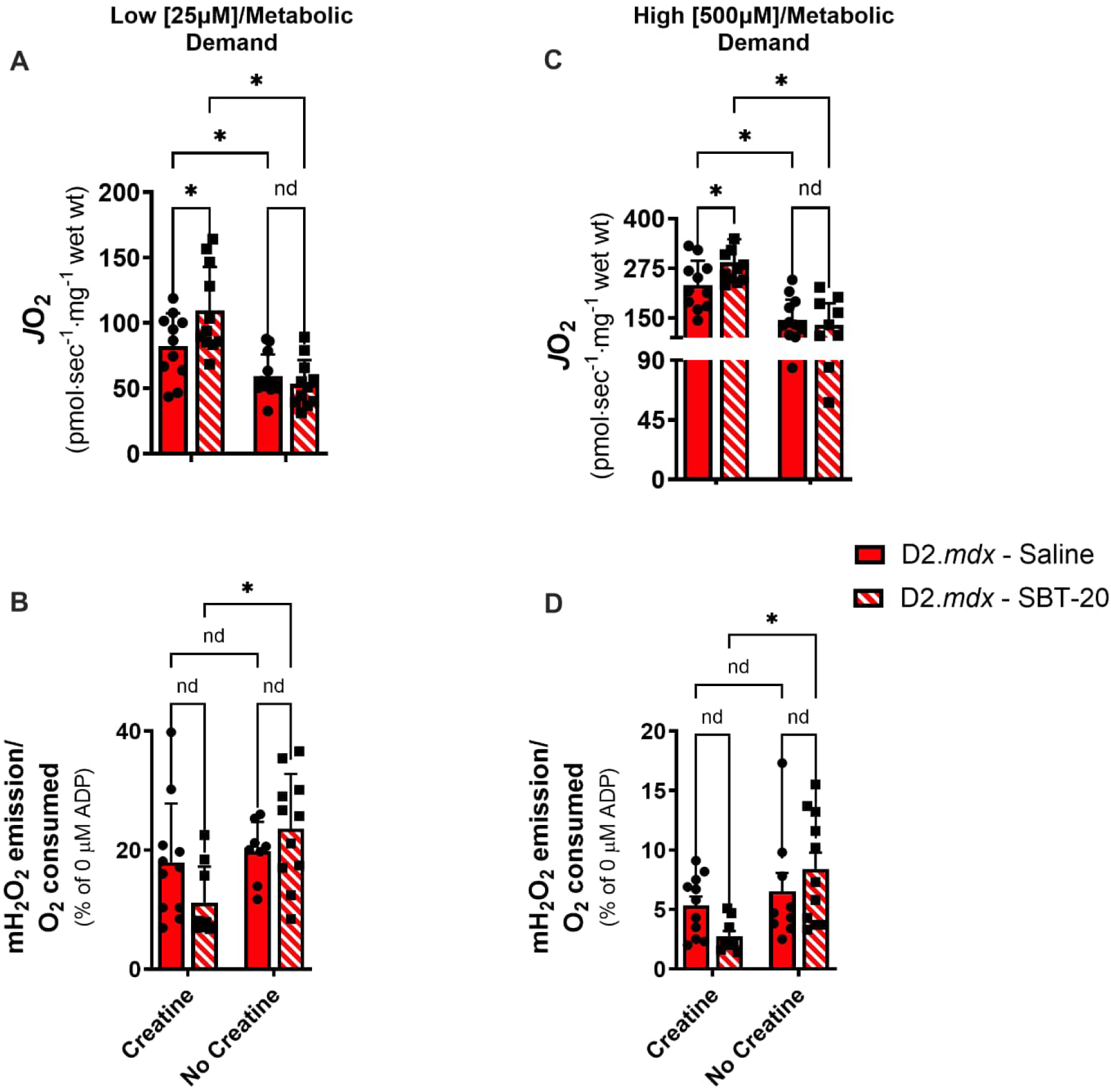
The effects of SBT-20 on complex I-supported mitochondrial respiration and mitochondrial H_2_O_2_ emission (mH_2_O_2_) in left ventricles from D2.mdx mice. Mice received daily subcutaneous injections of SBT-20 from day 4 to ∼12.5 weeks of age. Creatine-dependent and - independent ADP-stimulated respiration (JO_2_) and ADP-suppression of mH_2_O_2_ during the process of oxidative phosphorylation (mH_2_O_2_/O_2_) with low [ADP] (25uM, **A, B**) and high [ADP] (500uM; **C, D**) were assessed in cardiac left ventricle permeabilized muscle fibre bundles by stimulation with pyruvate (5mM respiration, 10mM mH_2_O_2_) and malate (2mM) to stimulate Complex I with NADH. Data were analyzed Two-way ANOVA for data in panels C and D Results represent mean +/- SD; n=8-12. *p<0.05; nd means ‘no difference’.

mH_2_O_2_ in the presence of creatine was not different than in the absence of creatine in D2.*mdx* saline treated mice (**Figure 4B and D**). Unlike the respiration data, this suggests that creatine insensitivity developed in D2.*mdx* at least in regard to regulating mH_2_O_2_. While a wildtype 12- week-old group was not included to verify this observation, it is well-established that creatine enhances the effect of ADP on attenuating mH_2_O_2_ (see discussion) as was seen in older 12-month wildtype mice (**Figure 2C and D**). This impaired ability of creatine to attenuate mH_2_O_2_ was not seen in SBT-20 treated mice given mH_2_O_2_ was lower with creatine compared to in its absence in both low and high [ADP] conditions (**Figure 4B and D**) thereby demonstrating that SBT-20 enhances creatine—dependent suppression of mH_2_O_2_ in line with the greater creatine-dependent respiration. Lastly, while it is not possible to determine if the compensatory increase in creatine-independent bioenergetics seen in 12-month D2.*mdx* mice (**Figure 2C, D**) occurred in these 12- week-old mice (**Figure 4A-D**) without a 12-week-old wildtype control group, the enhanced action of creatine on respiration and mH_2_O_2_ with SBT-20 nonetheless demonstrates its ability to improve mitochondrial creatine sensitivity, particularly given D2.*mdx* mice that did not receive SBT-20 were clearly insensitive to creatine.

Increased susceptibility to calcium-induced mitochondrial permeability transition pore activity was observed given calcium retention capacity was reduced in 12-month-old D2.*mdx* mouse left ventricles (**Supplemental Figure S1C**). There was no effect of SBT-20 on this measure compared to saline-treated D2.*mdx* mice at ∼12.5 weeks of age (**Supplemental Figure S1D**).

Select cardiac functional parameters were not altered by SBT-20 compared to saline-treated D2.*mdx* mice (**Supplemental Figure S4A-D**).

Lastly, to determine if higher mH_2_O_2_ in the presence of creatine was unique to the ADP-sensitive regulation of ROS production, we also assessed Complex III- and pyruvate dehydrogenase complex (PDC)-supported mH_2_O_2_ in the absence of ADP. In 12-month-old D2.*mdx*, Complex III- supported mH_2_O_2_ was lower in white gastrocnemius whereas PDC-supported mH_2_O_2_ was lower in the left ventricle, diaphragm, and quadriceps (**Supplemental Figure S1E)**. SBT-20 had no effect on these pathways. While this lower mH_2_O_2_ in 12-month-old D2.*mdx* mice warrant further investigation into how the contents or post-translational regulation of these pathways are altered during dystrophin deficiency, these data suggest the loss of creatine-dependent bioenergetics during pyruvate oxidation (**Figure 2**) is a unique mechanism contributing to higher mH_2_O_2_ during attenuated oxidative phosphorylation in DMD.

## Discussion

Creatine enhances the ability of ADP to stimulate oxidative phosphorylation and attenuate H_2_O_2_ emission in mitochondria. Here, we show 12-month-old D2.*mdx* mice have lower creatine-dependent bioenergetics that are related to greater cysteine oxidation of mtCK. In contrast, creatine-independent bioenergetics were apparently enhanced which may represent a form of mitohormesis in response to chronic disease in dystrophin deficient mice. Moreover, 12 weeks of treatment with the ROS-lowering mitochondrial-targeting peptide SBT-20 (up to ∼12.5 weeks of age) increased creatine-dependent respiration and lowered mH_2_O_2_, particularly under states of high metabolic demand in the left ventricle. These results demonstrate a specific mechanism linking redox and metabolic stress in mitochondria arising from dystrophin mutations and serve as a direction for continued development of mitochondrial-targeted therapies designed to restore metabolic and redox balance in DMD.

At 12 months of age, the left ventricle of D2.*mdx* mice also demonstrated a surprising increase in creatine-independent respiration, which contrasts the decreases we previously reported in this pathway in 4-week-old mice [6, 7]. This observation suggests the slower mitochondrial ADP-ATP system may compensate for impairments in the faster creatine-dependent phosphate shuttling mechanism [9]. The greater abundance of cysteine oxidation on mtCK was consistent with higher rates of creatine-sensitive mH_2_O_2_ during the process of oxidative phosphorylation and the more oxidized cellular environment (lower GSH:GSSG). mtCK oxidation may be unique to advanced stages of this disease, given we previously reported no differences in 4-week D2.*mdx* mice, at least in skeletal muscle [17]. Further studies could examine the precise form of thiol modification that occurs in 12-month-old D2.*mdx* mice, such as glutathionylation, considering its emerging role in linking redox signaling to metabolic control [22, 23] and considering the shift in GSH:GSSG noted in the present study. Likewise, the degree to which mtCK thiol redox state was preserved by SBT- 20 could be considered given the positive effects noted in this part of the study that was otherwise limited by tissue availability. Further studies could assess the specific cystines that were oxidized given cysteine 278, which regulates mtCK activity, and C358, which may regulate mtCK tethering to the inner mitochondrial membrane, were previously shown to be redox sensitive [24]. ANT and VDAC thiol oxidation could also be assessed to explain the apparent compensatory increases in creatine-independent bioenergetics at late stages of disease given their protein contents did not change, although reconciling these measures with the divergent response of creatine-dependent and -independent systems may be challenging given both proteins are thought to be primary regulators in either system.

We were unable to assess cardiac function in 12-month-old D2.*mdx* mice due to their qualitatively frail nature. Furthermore, while we did not see robust differences in grip strength compared to age-matched wild type mice, we did note that the absolute values of grip strength in the 12 month old wild type mice are ∼50% of the values we have reported previously in this strain at 4 weeks of age [7, 17] suggesting an aging effect occurred in the control group. Nonetheless, the other parameters demonstrate a severe myopathy in the 12-month-old D2.*mdx* mice. Also, the effect of age on mitochondrial reprogramming in locomotor and respiratory muscle pathology compared to muscle dysfunction could also be considered, particularly in relation to the earlier remodeling seen in 4- week-old D2.*mdx* [6, 7, 17, 18].

SBT-20 is a small tetrapeptide with high cell-penetrating potential that accumulates on cardiolipin in the inner mitochondrial membrane similar to the mitochondrial-targeting peptide elamipretide (formerly SS-31) [25–27] that prevents cytochrome *c* peroxidase activity, preserves oxidative phosphorylation, and prevents increases in superoxide production in response to stressors [28, 29]. SBT-20 also preserved mitochondrial respiration in H_2_O_2_-treated cells and partially prevents cardiac infarct size in response to ischaemia reperfusion injury [13]. To our knowledge, this is the first study to report a unique creatine-specific preservation of bioenergetics by cardiolipin-targeting peptides. As mtCK is thought to be bound to cardiolipin [9], future studies could consider whether the age-related impairment in creatine-dependent reductions in respiration seen in the present study was due to altered cardiolipin tethering to mtCK, and whether the preserved creatine-dependent bioenergetics by SBT-20 preserved such interactions. As ANT and VDAC are also thought to be bound to cardiolipin, the potential for SBT-20 to regulate the system as a whole could be considered.

As an aged-match wild type control group (∼12 weeks of age ) was not included for the SBT-20- treated D2.*mdx* experiments, we are not able to prove if the creatine-stimulated increases in respiration seen in D2.*mdx* vehicle treated animals were blunted compared to wildtype. We have previously demonstrated that creatine-dependent respiration is lower in 4-week-old D2.*mdx* mouse left ventricles [6]. Therefore, as creatine-dependent respiration is lower at both 4 weeks and 12 months as seen in **Figure 2**, it is possible that similar reductions would exist at ∼12 weeks as well, but this would require the inclusion of an age-matched wildtype control group to be certain. However, the results clearly demonstrate that creatine does not lower cardiac mH_2_O_2_ in 12-week-old D2.*mdx* vehicle-treated mice which is consistent with the creatine insensitivity seen in 4-week-old [6, 17] and 12-month-old D2.*mdx* hearts (**Figure 2C, D**) depending on the [ADP]. SBT-20 treatment lowered mH2O2 only in the presence of creatine which demonstrates a remarkable ability to convert mitochondria from a creatine-insensitive to creatine-sensitive phenotype in D2.*mdx* mice. Likewise, SBT-20 increased creatine-dependent respiration in D2.*mdx* mice to levels higher than what was seen in vehicle treated mice. As such, this study provides first-time proof-of-principle evidence that mitochondrial creatine-dependent bioenergetics can be enhanced by this class of mitochondrial-targeted therapeutics.

Although select cardiac functional parameters were not altered by SBT-20 compared to saline-treated D2.*mdx* mice, we cannot determine if a dysfunction existed at this age in comparison to wild type. In fact, prior work at younger ages in the D2.*mdx* mouse have reported no overt cardiomyopathy [6] while much older ages are known to have a moderate left ventricular cardiomyopathy in this model [19], at least as assessed with non-invasive approaches.

SBT-20 did not alter calcium retention capacity suggesting that it did not have an effect on mitochondrial permeability transition – an event that links mitochondrial calcium overload to apoptosis [30]. This is in contrast to previous reports showing SBT-20 prevents cytochrome *c* release, a key trigger of mitochondrial-induced apoptosis, in response to ischaemic stress [29] that is known to trigger mitochondrial permeability transition [30]. As our pilot study did not include a wild type control group, we are unable to determine whether calcium retention capacity was reduced in the D2.*mdx* saline control group at this age in contrast to the clear reductions seen at 12 months of age (**Supplemental Figure S1**).

Collectively, these findings suggest that a time-course design could be employed to determine whether SBT-20 prevents the unique time-dependent signatures of mitochondrial stress and delays the eventual onset of cardiomyopathy. To this end, we did not see an effect of SBT-20 on cardiac function assessed with echocardiography on ∼12.5-week-old mice. However, previous reports have shown an absence of overt cardiac dysfunction in D2.*mdx* mice at 4 weeks [6] or 7 weeks [19] of age but is apparent at 28 weeks and 52 weeks [19], with the latter age corresponding to our observation of oxidized mtCK. More in-depth analyses with invasive hemodynamics is also warranted given this approach could identify dysfunctions that are not detected with echocardiography.

### Additional Perspectives and Limitations

This investigation was designed to determine whether creatine-dependent and -independent mechanisms of phosphate shuttling remodel in positive or negative manners during advanced stages of disease in dystrophin deficient mice. The elevated mH_2_O_2_ was linked to a greater degree of oxidized cysteines in mtCK in 12-month-old D2.*mdx* mice. The use of a thiol labeling technique in immunoprecipitated mtCK provides a unique insight into a potential target of mitochondrial ROS during dystrophin deficiency that is linked to a specific impairment in creatine-dependent respiration. Thiol labeling of enriched protein fractions enabled this discovery that would not be possible if the study relied solely on broader measures of redox conditions in the cell. Rather, the change in glutathione redox state provides insight into how changes in mH_2_O_2_ impacted this primary H_2_O_2_ scavenger of the cell. Additional measures of lipid peroxidation or broader protein redox states in the cell could be considered in future investigations but are tangential to the question of this investigation pertaining to creatine-dependent and -independent bioenergetics. In the 2^nd^ investigation examining the effects of SBT-20, no redox measurements were performed to determine the impact of altered mH_2_O_2_ on cellular glutathione or other redox conditions or mtCK redox state. Such measurements should be considered in future investigations developing mitochondrial therapeutics for DMD in relation to measures of muscle function in order to more fully appreciate the impact of altered mitochondrial bioenergetics on redox-dependent processes regulating muscle contraction. Also, measurements of creatine, phosphocreatine, ADP and ATP (as examples) would add further insight into the degree to which the mitochondrial-cytoplasmic phosphate shuttling was altered in the cell. The extremely small muscle samples available from these 12-month-old frail dystrophin deficient mice were used in the most efficient way possible to obtain the dataset reported in this investigation such that future studies could consider adding these broader insights where possible.

## Conclusions

These findings demonstrate that left ventricle mitochondrial creatine metabolism is attenuated in relation to oxidized mitochondrial creatine kinase in late stages of disease in 12-month-old D2.*md*x mice. This impairment was linked to increases in creatine-independent bioenergetics which may represent a form of mitohormeses in response to chronic disease progression in dystrophin deficient mice. The ability of the ROS-lowering mitochondrial peptide SBT-20 to increase creatine-dependent pyruvate oxidation and lower creatine-dependent mitochondrial H_2_O_2_ emission demonstrates the potential for a mitochondrial-targeted therapeutic to enhance coupled respiration during dystrophin deficiency, particularly in regard to the regulation of mitochondrial creatine metabolism. This finding supports continued development of a new paradigm of mitochondrial- targeted redox and metabolic enhancing therapeutics that do not exist in the current standard of care of anti-inflammatory and other emerging treatments.

## Acknowledgements

We thank Trevor Teich for providing technical assistance with certain experiments, and Dr Robert Tsushima for kindly providing access to the Vevo 2100 ultrasound imaging system for echocardiography.

## Supplemental Information

Supplemental methods (Appendix A) and data (Appendix B) can be found in the separate file

*Supplemental Information*.

## Author Contributions

M.H., S.V.R., P.B., J.S. and C.G.R.P. contributed to the conception or design of the work. M.H., S.V.R., A.B., P.C.T., N.P., U.S., T.H., P.B., J.S. and C.G.R.P. contributed to acquisition, analysis and/or interpretation of data. All authors contributed to drafting the manuscript and approved the final version of the manuscript.

## Abbreviations

ADP: adenosine diphosphate
ATP: adenosine triphosphate
ANT: adenine nucleotide translocase
cCK: cytosolic creatine kinase
mH_2_O_2_: mitochondrial H_2_O_2_
mtCK: mitochondrial creatine kinase
PCr: phosphocreatine
PDC: pyruvate dehydrogenase complex
VDAC: voltage dependent anion carrier

