## Supplemental methods and figures for "Mitohormesis during advanced stages of Duchenne muscular dystrophy reveals a redox-sensitive creatine pathway that can be enhanced by the mitochondrial-targeting peptide SBT-20"

### Supplemental Information

#### Appendix A: Methodological Descriptions

Mice were provided access to standard chow and water ad libitum. All experiments and procedures were approved by the Animal Care Committee at York University (AUP Approval Number 2016-18) in accordance with the Canadian Council on Animal Care.

*SBT-20 injections and D2.mdx mice:*

D2.*mdx* mice receiving SBT-20 injections at day 4 of age were sexed at 10 days of age. At this point, treatment continued for 12 weeks only in male mice. Age-matched male D2.*mdx* mice received the volume equivalent dose of vehicle (0.9% NaCl; Saline) through subcutaneous injection. Half of the animals receiving 12 weeks of daily SBT-20 injections were allocated for cardiac function analysis through echocardiography as described previously [1] and the remaining animals were allocated for respiratory function analysis through esophageal pressure transduction [1] for data to be included in a separate manuscript. Animals were allowed to recover for 2 days following their respective test. On the day of sacrifice, prior to tissue harvest, animals underwent a μCT scan for muscle volume followed by *in vivo* hindlimb plantar flexor force production measurements [1] in a small subset of animals for data to be used in a separate manuscript.

*Voluntary Wheel Running*

Animals were placed in individual cages equipped with a locked 14 cm diameter running wheel and rotation counter (VDO m3 bike computer, Mountain Equipment Co-Op, Vancouver, Canada). After 24 hours of acclimatization, wheels were unlocked, and distance run over 24 hours was recorded as we have reported previously [1].

*Cage Hang Time*

Animals were placed on top of a metal cage lid and positioned so that all four limbs grasped the cage. The cage lid was inverted so that the mouse was hanging and cage hang time was recorded for a maximum of 180 seconds as previously described [2].

*Forelimb Grip Strength*

Forelimb grip strength was assessed using a force transducer (Mark 10 Digital Force Gauge, Copiague, NY) and has been described in detail previously [2]. Briefly, mice were removed from their cages by the tail and brought towards a grid attached to the force transducer until such time that the mice grasped the grid with their forepaws. Upon grasping, animals were pulled away from the grid until their grasp was broken. Peak tension was recorded, and the trial was repeated twice more. If the animal did not show resistance to the experimenter, the trial was not recorded. Maximum peak tension from the best of 3 trials was used for analysis [3].

*In Vivo μCT Scans*

*In vivo* body scans were performed using μCT (SkyScan 1278, Bruker-microCT, Kontick, Belgium) as previously described [2]. Following scans, lower limb muscle volume was analyzed through reconstruction of the image using NRecon (software version 1.7.0.4, Bruker-microCT, Kontick, Belgium) followed by the analysis of a selected region of interest using CTAn (software version 1.15.4.0, Bruker-microCT, Kontick, Belgium). The region of interest for hind-limb muscle volume was landmarked from the top of the patella to the ankle joint and was quantified by setting thresholds for fat free mass (76-102), expressed as volume ($mm^3$) of lean mass/g body weight.

*Preparation of Permeabilized Muscle Fibre Bundles (PmFB)*

This technique was performed in the same manner as we have reported previously [1, 4, 5]. Briefly, muscles were removed from mice during anaesthesia with isoflurane. Heart, diaphragm, quadriceps (Quad) and white gastrocnemius (WG) muscles were removed and immediately placed in ice-cold BIOPS buffer (pH 7.1). Each muscle was trimmed of connective tissue and fat and divided into several small muscle bundles (~2–7 mm, 1.0–2.5 mg wet weight). Each bundle was gently separated along the longitudinal axis to form bundles that were treated with 40 µg/ml saponin in BIOPS on a rotor for 30 minutes at 4° C. Bundles for pyruvate-stimulated Complex I- and PDC-supported mitochondrial $H_2O_2$ emission ($mH_2O_2$) measurements (described below) were also treated with 35 µM 2,4-dinitrochlorobenzene (CDNB) during the permeabilization step to deplete glutathione and allow for detectable rates of $mH_2O_2$ [6]. Following permeabilization, the PmFB were divided into three groups; 1) bundles intended for high resolution respirometry were placed in MiR05 (pH 7.1) buffer while 2) bundles intended for $mH_2O_2$ were placed in Buffer Z (pH 7.4). PmFB were washed on a rotor at 4°C in MiR05 or Buffer Z (<4 hours) until the measurements were initiated. Following permeabilization and washing, all mitochondrial bioenergetic measurements were performed in the same order from one animal to another and between groups to ensure consistent durations of the wash step.

*High-resolution respirometry*

High-resolution $O_2$ consumption measurements were conducted in 2 ml of respiration medium (MiR05) using the Oroboros Oxygraph-2k (Oroboros Instruments, Corp., Innsbruck, Austria) with stirring at 750 rpm at 37 °C [7-11]. Respiration medium contained 20mM Cr to saturate mitochondrial creatine kinase or no creatine to prevent the activation of mtCK [9, 12]. A third condition of 13.9mM PCr and 9.1mM Cr was used to provide an equilibrium across mtCK as occurs in skeletal muscle at rest *in vivo* [13] noting we were unable to find concentrations for contracting cardiac muscle. For ADP-stimulated respiratory kinetics, 5 mM pyruvate, accompanied by 2 mM malate, were added as Complex I substrates (via generation of NADH to saturate electron entry into Complex I) followed by a titration of physiological ADP (25 µM) and supraphysiological (500 µM) ADP to mimic low and high metabolic stress respectively. Cytochrome *c* was added to test for mitochondrial membrane integrity, with all experiments demonstrating <10% increase in respiration. All experiments were conducted in the presence of 5µM BLEB in the assay media to prevent spontaneous contraction of PmFB [9, 10, 14]. Each protocol was initiated with a starting [$O_2$] of approximately 350 µM and was completed before the oxygraph chamber [$O_2$] reached 150 µM as done previously [7-11]. Polarographic oxygen measurements were acquired in 2 s intervals with the rate of respiration derived from 40 data points and expressed as pmol/s/mg wet weight. PmFB were weighed in ~1.5 ml of tared BIOPS (ATP-containing relaxing media) to prevent rigor.

*Mitochondrial $H_2O_2$ Emission ($mH_2O_2$)*

$mH_2O_2$ was determined fluorometrically (QuantaMaster 40, HORIBA Scientific, Edison, NJ,

USA) in a quartz cuvette with continuous stirring at 37°C, in 1 mL of Buffer Z supplemented with 10 μM Amplex Ultra Red, 0.5 U/ml horseradish peroxidase, 1 mM EGTA, 40 U/ml Cu/Zn-SOD1, 5 μM BLEB and 20mM Cr. Site specific induction of $mH_2O_2$ was measured through the addition of either 10 mM pyruvate and 2 mM malate (NADH, Complex I), 10 mM succinate ($FADH_2$, Complex I via reverse electron flux from Complex II) or 2.5 μM antimycin A (Complex III). Additionally, using 0.5 μM rotenone, a complex I inhibitor, plus 10 mM pyruvate, electron slip specific to pyruvate dehydrogenase complex was also measured in CDNB-treated fibres as noted above [6]. Following the induction of state II $mH_2O_2$ by Complex I and Complex II substrates, a titration of ADP was added to progressively attenuate $mH_2O_2$. Complexes I and II-supported $mH_2O_2$ were repeated with no creatine in the assay buffer to compare ADP's effects without mtCK-mediated phosphate shuttling. All measurements were made in the presence of 1 μM BLEB to prevent ADP-induced rigor as described above. After the experiments, the fibres dried and weighed as described above. The rate of $H_2O_2$ emission was calculated from the slope (F/min), from a standard curve established with the same reaction conditions and normalized to fibre bundle dry weight.

*Glutathione*

Glutathione was assessed as previously described [15]. GSH was assessed by UV-HPLC monitoring of NEM-GSH while GSSG was assessed by fluorescent-HPLC by tracking O-pthalymide (OPA, Sigma-Aldrich, Oakville, Canada) tagged GSH through a flow-through cuvette following GSSG conversion to GSH (FireflySci 8830, NY, USA) in a QuantaMaster 40 spectrofluorometer (HORIBA, NJ, USA). All values were normalized to protein concentration and reported in $\mu$mol/g protein. All measures were performed in lysate prepared from the entire heart remaining after left ventricles were removed for permeabilized fibre preparation described above.

*Western Blotting*

After removing left ventricles for preparing permeabilized fibres, the remaining heart was frozen in liquid nitrogen. A piece of this sample as well as a piece of frozen diaphragm, quadriceps and white gastrocnemius (10–30 mg) from each mouse were homogenized in a plastic microcentrifuge tube with a tapered teflon pestle in ice-cold buffer containing (mm): 40 HEPES, 120 NaCl, 1 EDTA, 10 $NaHP_2O_7 \cdot 10H_2O$ pyrophosphate, 10 β-glycerophosphate, 10 NaF and 0.3% CHAPS detergent (pH 7.1 adjusted using KOH). Protein concentrations were determined using a BCA assay (Life Technologies, Carlsbad, CA, USA). Fifty μg of denatured and reduced protein was subjected to 10–12% gradient SDS-PAGE followed by transfer to low-fluorescence polyvinylidene difluoride membrane. Membranes were blocked with LI-COR Odyssey Blocking Buffer (LI-COR, Lincoln NE, USA) and immunoblotted overnight (4°C) with antibodies specific for each protein. A commercially available monoclonal antibody was used to detect electron transport chain proteins (human OXPHOS Cocktail, ab110411; Abcam, Cambridge, UK, 1:250 dilution), including V-ATP5A (55 kDa), III-UQCRC2 (48 kDa), IV-MTCO1 (40 kDa), II-SDHB (30 kDa) and I-NDUFB8 (20 kDa). Commercially available polyclonal antibodies were used to detect voltage dependent anion carrier 2 (VDAC 2) (32059, 32 kDa; Santa-Cruz, 1:1000), adenine nucleotide translocase 1 (ANT 1) (ab180715, 33 kDa; Abcam, 1:1000). An antibody for sarcomeric s-mtCK

(42 kDa, 1:1000) and purified mtCK were generous gifts from Dr Uwe Schlattner, Grenoble, France. The mtCK antibody has been validated previously to confirm specificity [16].

After overnight incubation in primary antibodies, membranes were washed three times, for 5 minutes each time, in TBST and incubated for 1 hour at room temperature with the corresponding infrared fluorescent secondary antibody (LI-COR Biotechnology, Lincoln, NE, USA). Immunoreactive proteins were detected by infrared imaging (LI-COR CLx; LI-COR Biotechnology, Lincoln, NE, USA) and quantified by densitometry (ImageJ, http://imagej.nih.gov/ij/). All images were normalized to a whole membrane Amido Black total protein stain (A8181, Sigma, St Louis, MO, USA).

*Redox Status of MtCK*

Free reactive thiol (SH) groups on mtCK were labeled using IR-Dye 800CW-Maleimide (LiCor Biotechnology, Lincoln, NE, USA) by a modification of the technique previously described [17, 18]. A portion of frozen heart (after left ventricles were removed) from 12-month-old mice was homogenized in CHAPS buffer plus inhibitors as described above. Samples were incubated overnight at 4°C with the IR-Dye at a concentration of 100nM/200µg total protein. Excess dye was removed from labeled proteins over Zeba Desalting Spin Columns (Thermo Fisher Scientific, Burlington, Canada) and protein concentration was determined using a BCA assay (Life Technologies, Carlsbad, CA, USA). MtCK was incubated at a ratio of 1:50 with 1mg of SureBead Protein G Magnetic Beads (Biorad, Mississauga, Canada) for 10 minutes at room temperature. 600µg of labeled proteins were then incubated with the SureBeads for 1 hour at room temperature. The bound proteins were eluted with 20mM Glyicine (pH 2.0) and were subjected to 10% SDS-PAGE. The gels were scanned using infrared imaging (LI-COR CLx; LI-COR Biotechnology, Lincoln, NE, USA) and quantified by densitometry (ImageJ, http://imagej.nih.gov/ij/). IR-dye 800 CW fluorescence was normalized to total mtCK from a portion of left ventricles taken from the same mice in a separate blot.

*Statistics*

Results are expressed as means ± SD. The level of significance was established as $p<0.05$ for all statistics. Outliers were omitted in accordance with the ROUT test GraphPad Prism Software, La Jolla, CA, USA). D'Agostino-Pearson omnibus normality test revealed that data resembled a Gaussian distribution which justified the application of parametric tests. Statistical differences were analyzed using Two-way ANOVA with or unpaired t-tests as indicated in figure legends. When significant interactions were observed with ANOVA, post-hoc analyses of multiple group comparisons were performed using a two-stage set-up method of Benjamini, Krieger and Yekutieli to control the false discovery rate (FDR) using GraphPad Prism 10 (La Jolla, CA, USA).

**Appendix B: Supplemental Figures**

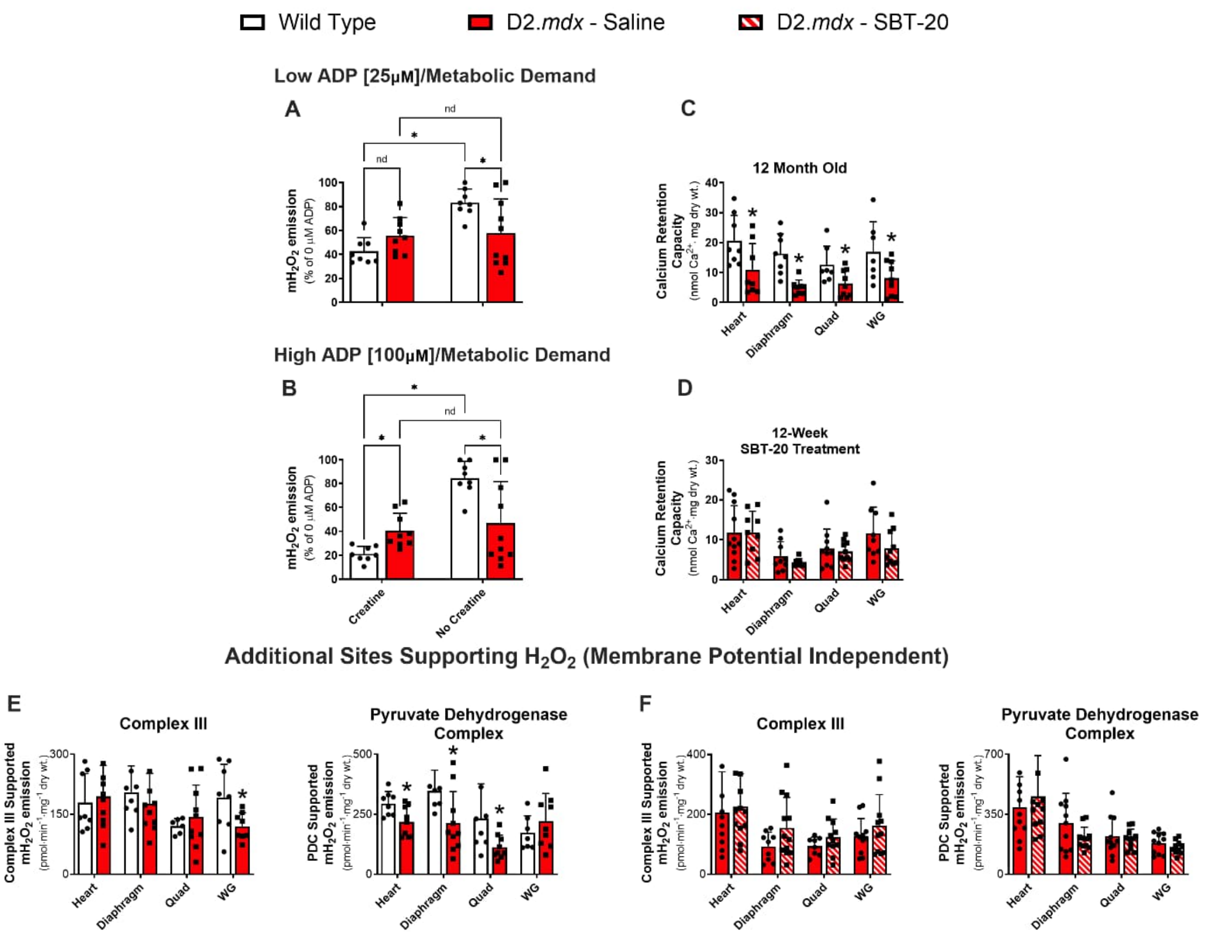


**Supplemental Figure S1. Additional sites supporting $mH_2O_2$ and mitochondrial calcium retention capacity in 12-month-old or SBT-20-treated D2.*mdx* mice.**

In 12-month-old wildtype and D2.*mdx* mice, Complex II-supported $mH_2O_2$ in the presence of low [ADP] (25uM; **A**) and high [ADP] (500uM; **B**), with and without creatine in the media, was achieved with 10mM succinate ($FADH_2$) which causes superoxide production at Complex I due to reverse electron transfer that is subsequently dismutated to $H_2O_2$ by endogenous matrix superoxide dismutase. Mitochondrial calcium retention capacity was assessed in muscles as described in A and B in 12-month-old wildtype and D2.*mdx* mice (**C**) and mice receiving daily saline or SBT-20 subcutaneous injections from day 4 to ~12.5 weeks of age (**D**). $mH_2O_2$ arising from Complex III stimulation by 2.5μM antimycin A and from pyruvate dehydrogenase complex (PDC) stimulation by 10mM pyruvate and 0.5μM rotenone (to block NADH oxidation by Complex I forcing reduction of PDC) were assessed in permeabilized fibres from each muscle (**E, F**). Measures were performed in permeabilized fibres from cardiac left ventricles (Heart), diaphragm, quadriceps (Quad) and white gastrocnemius (WG). Data were analyzed by *Two-way ANOVA for data in panels A and B, and unpaired t*-tests in remaining data. Results represent mean ± SD; n=6-12. *p<0.05 compared with saline-treated D2.*mdx* mice within the same “Creatine” or “No Creatine” condition (A, B) or muscle type (C-F); *nd means ‘no difference’*.

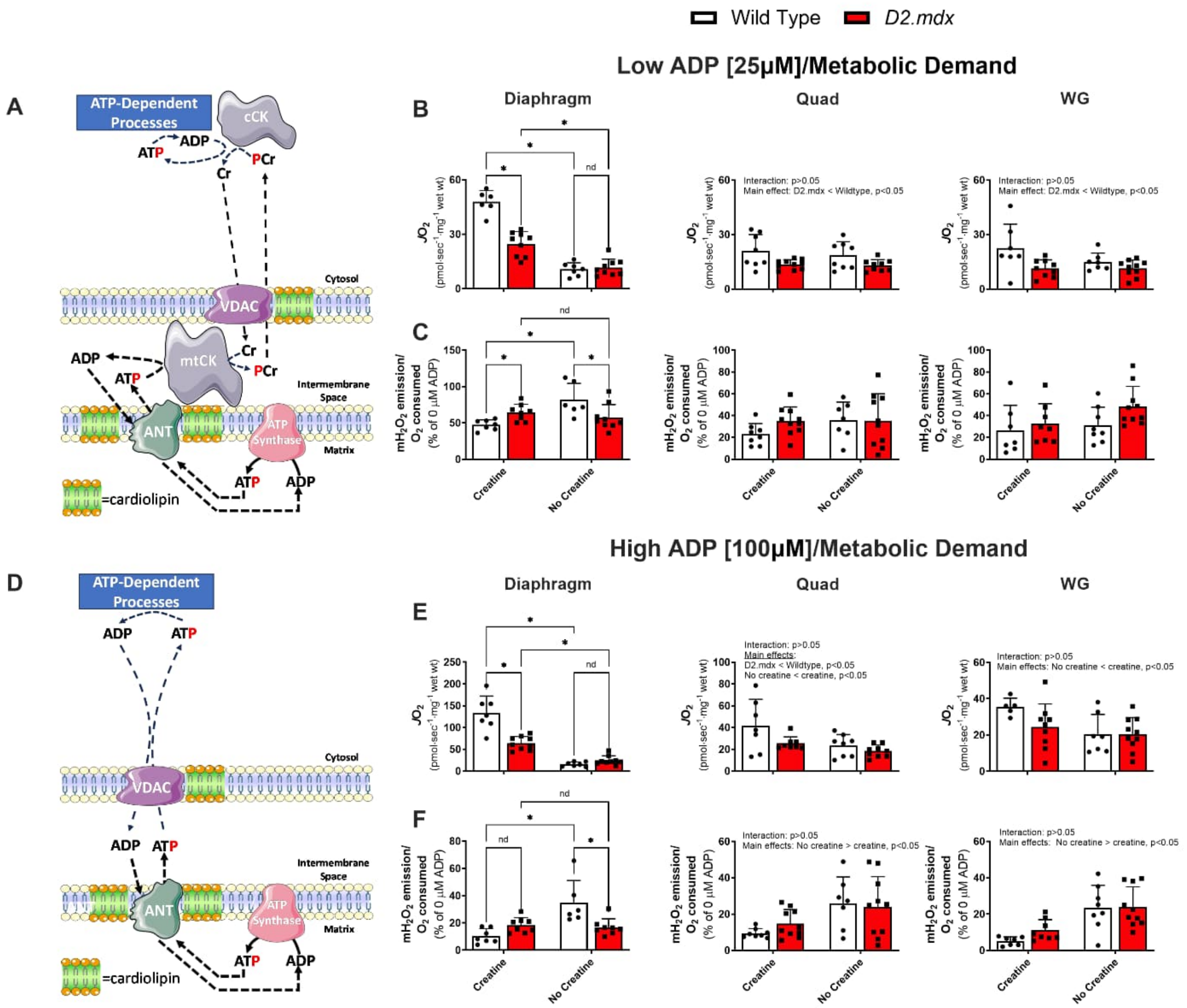


**Supplemental Figure S2**. **Complex I-supported mitochondrial respiration and mitochondrial $H_2O_2$ emission ($mH_2O_2$) in diaphragm, quadriceps (Quad) and white gastrocnemius (WG) from 12-month-old D2.*mdx* mice.**

Creatine-dependent (**A**) and -independent (**B**) ADP-stimulated respiration ($JO_2$) and ADP-suppression of $mH_2O_2$ during oxidative phosphorylation ($mH_2O_2/O_2$) with low [ADP] (25uM; **C, D**) and high [ADP] (500uM; **E, F**) were assessed in permeabilized muscle fibre bundles by stimulation with pyruvate (5mM for respiration experiments, 10mM for $mH_2O_2$ experiments) and malate (2mM) to stimulate Complex I with NADH. (See **Figure 1** in the manuscript for more detailed description of creatine-dependent and -independent phosphate shuttling (A,B) and the governance of $mH_2O_2$). $mH_2O_2/O_2$ was expressed as a % of maximal $H_2O_2$ achieved with pyruvate malate alone prior to the addition of ADP in the titration protocol (data not shown). Data were analyzed by t-tests between wildtype and D2.*mdx*. Results represent mean ± SD; n=5-10. *p<0.05 compared with wildtype within the same “Creatine” or “No Creatine” condition.

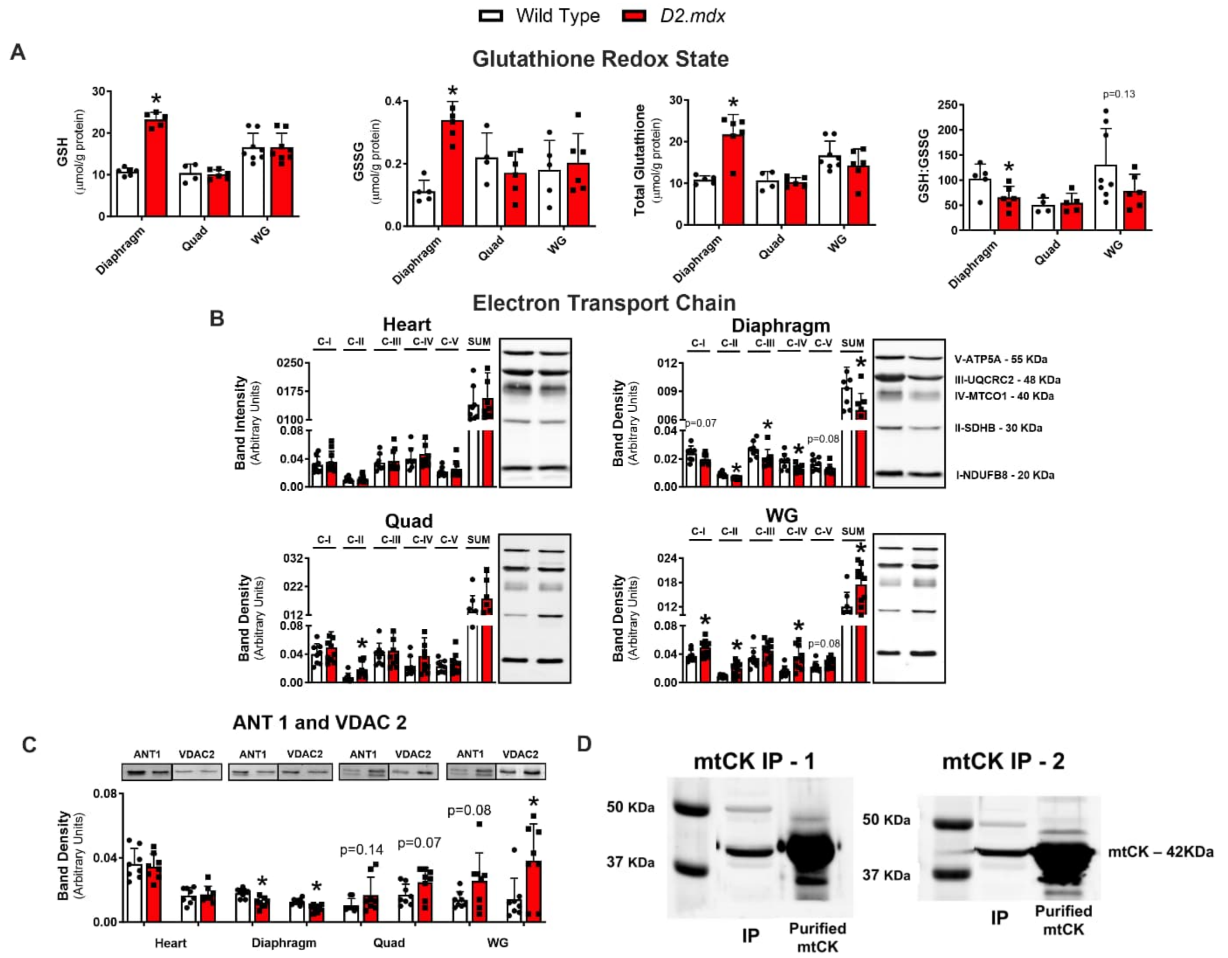


**Supplemental Figure S3**. **Cellular glutathione equilibrium, protein contents of electron transport chain components, ANT1, VDAC2, and mtCK immunoprecipitation validations.**

Glutathione redox state (**A**) was measured in skeletal muscle lysates using HPLC-UV for the detection of GSH and HPLC-fluorescence GSSG (**B**). The GSH:GSSG ratio and total glutathione (GSH + 2x GSSG) were calculated from GSH and GSSG. All glutathione measures were made in diaphragm, quadriceps (Quad) and white gastrocnemius (WG). Western blot-detection (**B, C**) of total protein contents of mitochondrial adenine nucleotide translocase (ANT) 1, voltage dependent anion carrier (VDAC) 2 and specific electron transport chain subunits of Complexes I-IV (C-I, C-II, C-III, C-IV) and ATP synthase (CV), and their sum, in lysates from frozen heart (following removal of left ventricles), diaphragm, Quad and WG. Verification of banding in immunoprecipitated mtCK from heart (following removal of left ventricles) relative to purified mtCK standards (**D**). All data were analyzed by unpaired *t*-tests between wildtype and D2.*mdx*. Results represent mean ± SD; n=4-10. *p<0.05 compared with wildtype.

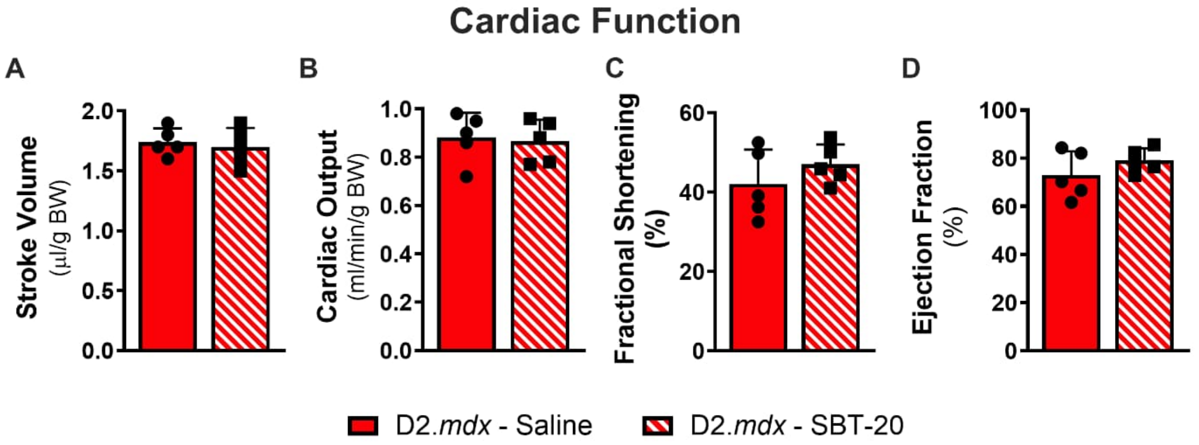


**Supplemental Figure S4**. **Assessments of cardiac function by echocardiography.**

D2.*mdx* mice received daily subcutaneous injections of SBT-20 from day 4 to ~12.5 weeks of age. Cardiac functional measures of stroke volume (**A**), cardiac output (**B**), fractional shortening (**C**) and ejection fraction (**D**) were assessed with echocardiography in saline- and SBT-20-treated D2.*mdx* mice. Data were analyzed by *t*-tests between wildtype and D2.*mdx*. Results represent mean ± SD; n=5 for cardiac function.
